# Social Interaction Elicits Activity in Glutamatergic Neurons in the Posterior Intralaminar Complex of the Thalamus

**DOI:** 10.1101/2023.04.24.538114

**Authors:** Leithead A.B., Godino A., Barbier M., Harony-Nicolas H.

## Abstract

**Background:** The posterior intralaminar (PIL) complex of the thalamus is a multimodal nucleus that has been implicated in maternal behaviors and conspecific social behaviors in male and female rodents. Glutamatergic neurons are a major component of the PIL; however, their specific activity and role during social interactions has not yet been assessed.

**Methods:** We used immunohistochemistry for the immediate early gene c-fos as a proxy for neuronal activity in the PIL of mice exposed to a novel social stimulus, a novel object stimulus, or no stimulus. We then used fiber photometry to record neural activity of glutamatergic neurons in the PIL in real-time during social and non-social interactions. Finally, we used inhibitory DREADDs in glutamatergic PIL neurons and tested social preference and social habituation-dishabituation.

**Results:** We observed significantly more *c-fos*-positive cells in the PIL of mice exposed to social versus object or no stimuli. Neural activity of PIL glutamatergic neurons was increased when male and female mice were engaged in social interaction with a same-sex juvenile or opposite-sex adult, but not a toy mouse. Neural activity positively correlated with social investigation bout length and negatively correlated with chronological order of bouts. Social preference was unaffected by inhibition; however, inhibiting activity of glutamatergic neurons in the PIL delayed the time it took female mice to form social habituation.

**Conclusions:** Together these findings suggest that glutamatergic PIL neurons respond to social stimuli in both male and female mice and may regulate perceptual encoding of social information to facilitate recognition of social stimuli.

## INTRODUCTION

The posterior intralaminar complex of the thalamus (PIL) is considered a relay center by which sensory information is conveyed to other brain regions for the generation of relevant social behaviors (1–6). Tracing studies conducted in rodents have provided anatomical support for this theory, demonstrating that the PIL receives inputs from sensory regions such as the inferior and superior colliculus, auditory cortex, and spinal cord (3, 7–10) and projects to socially relevant brain regions including the paraventricular nucleus of the hypothalamus (PVH), the lateral septum, medial preoptic area (MPOA), and the medial amygdala (11).

Functional studies have also supported the role of the PIL as a relay center for sensory information. In maternal female rats, a subpopulation of PIL neurons that expresses tuberoinfundibular peptide 39 (TIP39) were shown to be activated in response to pup exposure and suckling (2–4). These neurons co-express calbindin and vesicular glutamate transporter 2 (VGlut2), and form presumed excitatory synapses with oxytocin neurons in the PVH (3, 12), which are known to regulate a range of social behaviors, from parental care to social recognition of conspecifics (13–18). The PIL has also been shown to be activated in rats following the reunion of conspecific females (3) and more recently, shown to be involved in grooming behavior in males and females (11). While together these findings suggest that the PIL may play a role in social behavior, it remains unknown how PIL neurons respond, in real-time, to social interactions with novel same-sex or opposite-sex counterparts. Additionally, the specific contribution of glutamatergic neurons to social behaviors has not yet been assessed.

We hypothesized that neural activity in glutamatergic PIL neurons is required and therefore increased during social interactions in both male and female mice. We also hypothesized that inactivation of PIL glutamatergic neurons would have a deleterious effect on social behavior. To address our hypotheses, we first used the immediate early gene *c-fos* to examine the activity of PIL neurons following social interaction with a novel same-sex juvenile in both male and female mice. We then employed fiber photometry calcium imaging to follow the activity of glutamatergic PIL neurons in behaving mice, during interaction with a novel same-sex juvenile, novel opposite-sex adult, and novel object stimulus (as a control). Finally, we used chemogenetic tools to examine the impact of inhibition of glutamatergic PIL neurons on social behavior, in tasks of social preference and social habituation-dishabituation.

## METHODS AND MATERIALS

For detailed methods relating to stereotaxic surgery, histology, immunohistochemistry, RNAscope, and fiber photometry analyses, please see Supplement.

### Animals

Adult (8-12 weeks) male and female C57BL/6 mice (Taconic Biosciences) were used as experimental subjects. Opposite-sex adults (8-12 weeks) and same-sex juveniles (4-5 weeks) of the same strain were used as social stimuli in behavioral assays. Animals were housed in groups of 3-5 mice under a 12-hour light/dark cycle at a temperature of 22 ± 2°C with food and water available *ad libitum*. Experiments were conducted during the light phase cycle. Procedures were carried out in accordance with protocols approved by the Institutional Animal Care and Use Committee at the Icahn School of Medicine at Mount Sinai.

### Viral vectors

For retrograde tracing experiments we used AAVRetro-hSyn-eGFP-Cre (Cat. #105540, Addgene). To capture glutamatergic neural activity, we used a 1:1 combination of AAV8-CaMKIIα-mCherry-Cre (University of North Carolina [UNC] Vector Core) and AAV9-CAG-Flex-GcaMP6m (Cat. #100839, Addgene). To inhibit glutamatergic neurons, we used a 1:1 combination of AAV8-CaMKIIα-GFP-Cre (UNC) and AAV2-hSyn-DIO-hM4D(Gi)-mCherry (Cat. #44362, Addgene).

### Drugs

For chemogenetic inhibition using DREADDs, clozapine-N-oxide (CNO) was dissolved in vehicle solution (0.05% DMSO + 0.9% saline) at a concentration of 1 mg/mL then injected intraperitoneally (IP; 3.5 mg/kg)(19).

### *c-fos* induction and immunohistochemistry

To quantify *c-fos* expression in the PIL in response to social and nonsocial stimuli, male and female adult mice were habituated to an open field box for 15 minutes then exposed to a novel same-sex juvenile, a novel object toy mouse, or no stimulus (n = 3/sex/condition). They were allowed to freely interact for 1 hour, then immediately after underwent perfusion and brain collection for *c-fos* staining using immunohistochemistry, which is detailed in the Supplement.

### Behavioral assays

#### Social/Object Preference

In the social/object preference task (20), either a social stimulus (same-sex juvenile or opposite-sex adult) or object (toy mouse) was placed into one compartment with the opposite compartment left empty (counterbalanced) and the subject mouse was allowed to interact for 10 minutes.

#### Social Habituation-Dishabituation

In the social habituation-dishabituation paradigm (21–24), a novel same-sex juvenile was placed into the box and free social interaction was permitted for 5 minutes (T1). The juvenile was then removed from the box, leaving the subject mouse alone in the box for a 10-minute break. This series was then repeated 3 more times with the same juvenile (T2-T4) to capture habituation to the stimulus. A different novel same-sex juvenile was then introduced for one final 5-minute bout (T5) to control for lack of motivation to engage in social interaction.

### Fiber Photometry Recording

To measure the bulk activity of glutamatergic PIL neurons during social interaction, male and female adult mice underwent 3 separate days of recording during social same-sex juvenile, social opposite-sex adult, and control toy mouse preference tasks (n = 9/sex). Behavior and calcium activity were recorded using methods previously described (25) with a 400-um 0.57 N.A. fiberoptic patchcord (Cat. #MFP_400/430/1100-0.57; Doric). More detailed information is available in the Supplement. Subjects were excluded from analyses based on lack of viral expression (n = 2 males) or fiber misplacement observed upon postmortem inspection of the injection site (n = 1 male, 2 females).

### DREADDs

To assess the impact of inhibiting PIL glutamatergic neurons on social behavior, male and female mice were bilaterally injected with inhibitory DREADDs in the PIL. Three weeks later, mice were handled and habituated to IP injections of vehicle solution. Mice were then tested in the social preference task (Cohort 1, n = 9 males, 8 females) and/or social habituation-dishabituation paradigm (Cohort 1 + 2, n = 17 males, 12 females) on separate testing days following IP injection of either CNO (3.5 mg/kg) or the vehicle solution of an equal volume (within-subjects, counterbalanced) 15 min before testing. Mice were excluded from analyses for low viral expression of DREADDs (n = 1 male, 3 females) or aggressive behavior during free social interaction in the habituation-dishabituation paradigm (n = 3 males).

### Analyses

Behavior, fiber photometry, and imaging analyses are detailed in the Supplement and Data Table. For each measure, differences between sex as a variable were assessed and when nonsignificant, data from both sexes are presented together. No statistical power estimation analyses were used to predetermine sample sizes, which instead were chosen to match previous publications (2, 11, 26-28).

## RESULTS

### Novel social interaction elicits *c-fos* activation in the PIL

Previous research in rats and mice has shown that neuronal activity in the PIL is increased during mother-pup interactions, as well as during social interaction between familiar female conspecifics (3, 11). However, whether neuronal activity is increased in males and females during an interaction with an unfamiliar social stimulus has not been investigated. To examine PIL neuronal activity in response to novel stimuli, we quantified the number of neurons that express the immediate early gene, *c-fos,* in the PIL as a proxy for neuronal activity, following a one-hour free social interaction between an adult subject mouse and an unfamiliar same-sex juvenile mouse, a non-social stimulus (toy mouse), or no stimulus (Fig. 1A-B). The number of *c-fos*^+^ cells in the PIL was higher in male and female mice exposed to a novel social stimulus (*M =* 485*, SD =* 64.64), compared to those exposed to a novel object stimulus (*M =* 385.1*, SD =* 41.42) or no stimulus (*M* = 233.2, *SD* = 51.52*; F*(2,11) = 28.52, *p* < .0001; Fig. 1C). These findings suggested that activity in the PIL is significantly increased in the presence of a social stimulus.

**Figure 1.**
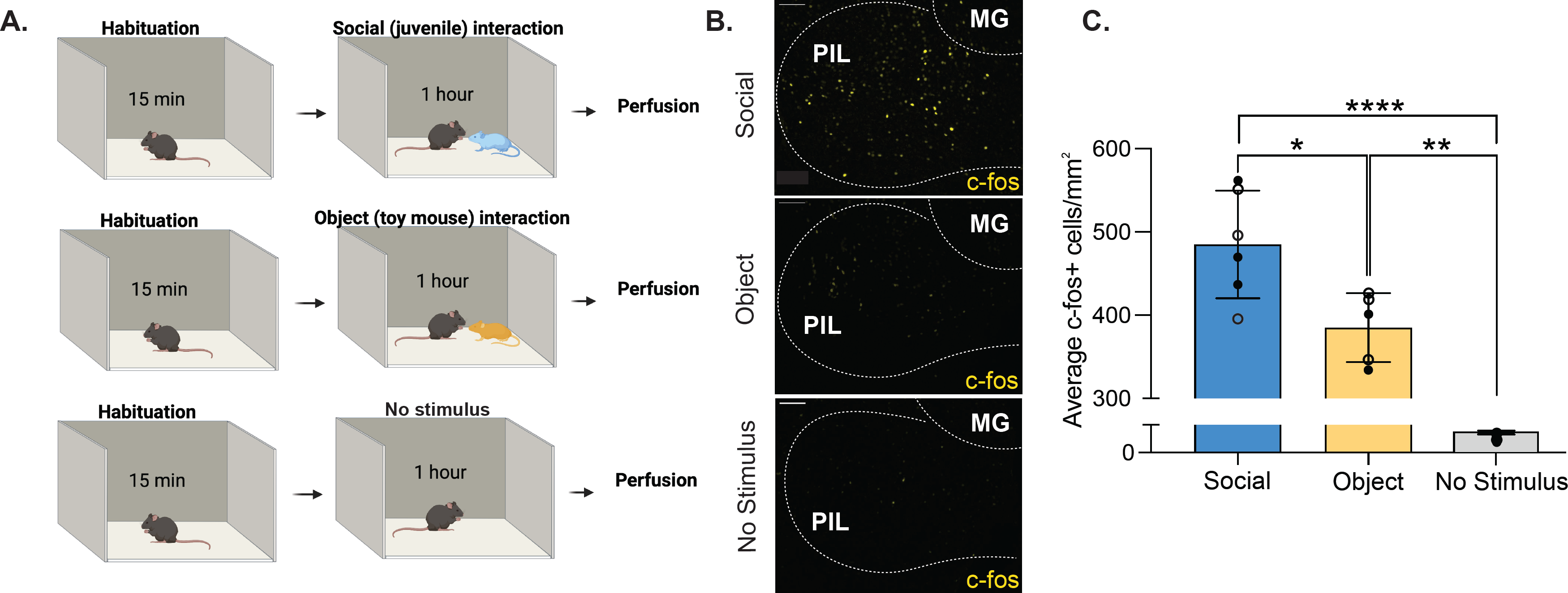
Social interaction elicits more *c-fos* expression in the PIL than object interaction or no stimulus interaction in males and females. **(A)** Schematic of social, object and no stimulus interaction paradigms. **(B)** Sample patterns of *c-fos* expression in the PIL using immunohistochemistry with anti-*c-fos* antibodies (yellow) in response to a social stimulus (upper panel), an object stimulus (middle panel), or no stimulus (lower panel). [MG, medial geniculate; 10X magnification; scale bar (100 μm)]. **(C)** Quantification of *c-fos^+^* cells per mm^2^ in the PIL of male and female mice reveal higher *c-fos* expression in response to a same-sex social stimulus compared to an object (toy mouse) or no stimulus. Each point represents one subject (13-15 sections/subject). Open symbols denote male subjects and closed symbols denote female subjects. (Two-way between-subjects ANOVA, *\*p<.*05, \*\**p*<.01, ****p<.0001; no sex difference; n = 3/sex/condition).

Previous research has suggested that excitatory projections from the PIL to the PVH are involved in social interactions in females and may contribute to oxytocin release via direct inputs to oxytocin cells (3). By combining retrograde tracing in the PVH and RNAscope in the PIL (using *vglut2* probe), we confirmed the existence of glutamatergic projections from the PIL to the PVH in mice (Supplementary Fig. 1). We therefore decided to further focus our studies on glutamatergic activity in the PIL.

### Real-time activity of glutamatergic PIL neurons increases during novel same-sex and opposite-sex social interactions, but not novel object interaction

We aimed to expand upon our *c-fos* findings by capturing real-time PIL activity in response to same-sex social stimuli, as well as opposite-sex social stimuli, while focusing on glutamatergic neurons. To gain temporal resolution of PIL neuronal activity, we used *in vivo* fiber photometry calcium imaging coupled with a genetically encoded calcium indicator (GcaMP). To specifically target glutamatergic PIL neurons, we used a dual viral approach where one virus expresses a cre-recombinase under the control of a CaMKIIα promoter, thus providing specificity to glutamatergic neurons (Supplementary Fig. 2), and the other expresses a cre-dependent GcaMP (Fig. 2A). Three weeks following viral expression, GcaMP6m fluorescent signals were recorded on three separate testing days: interaction with a novel same-sex juvenile, interaction with a novel opposite-sex adult, and interaction with a novel object (moving toy mouse). In all tests, the stimulus mouse or object was contained in one of two opposing compartments within the testing arena (Fig. 2B, upper panel), thus allowing the transmission of auditory, visual, and olfactory, but not somatosensory (touch) information.

**Figure 2.**
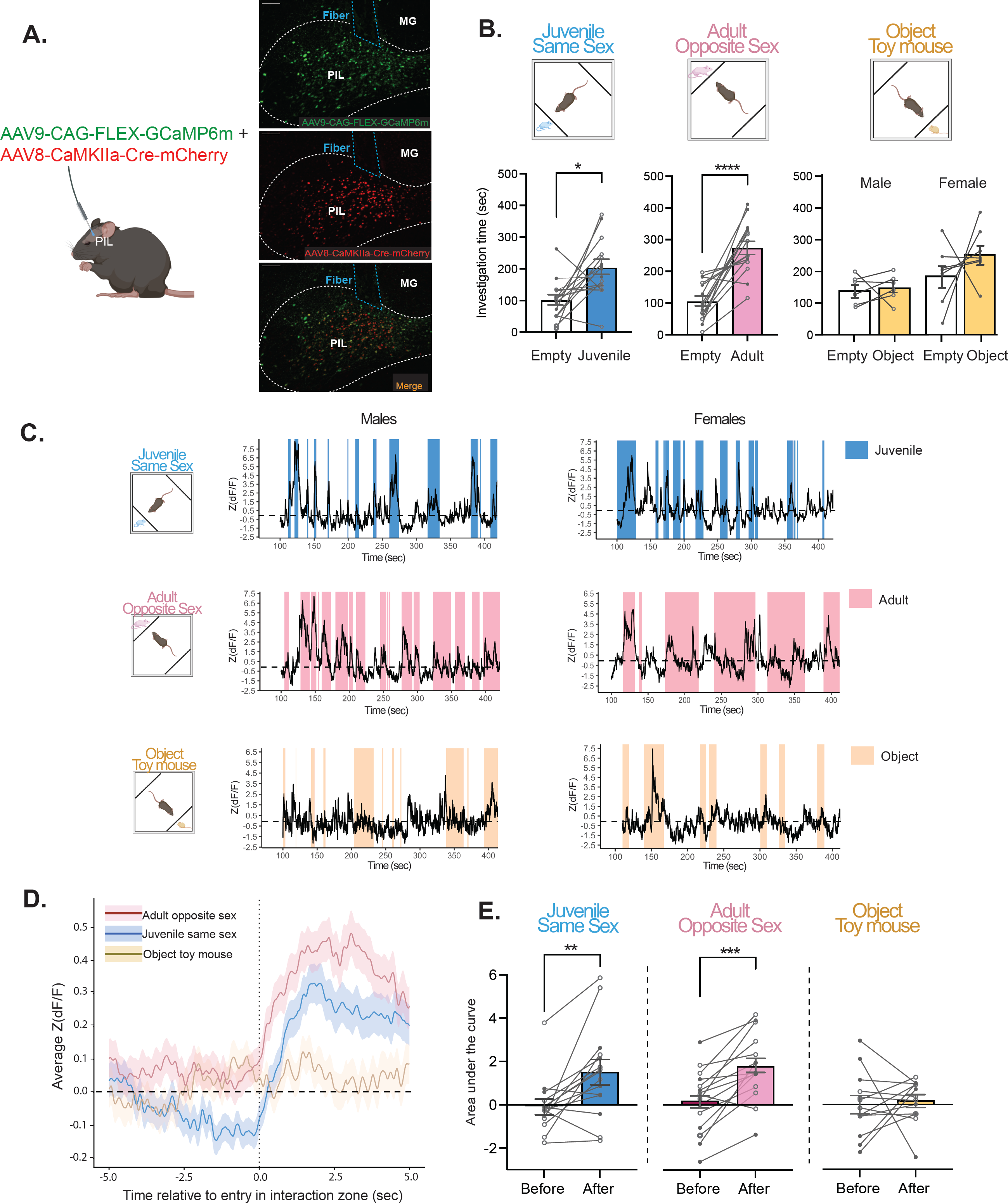
Activity of PIL glutamatergic neurons is increased during same-sex and opposite-sex social interactions, but not object interaction, in males and females. **(A)** Schematic of viral injection and fiber photometry, with sample histology showing GCaMP6m expression (green; top), CaMKIIα-Cre-mCherry (red; middle), and overlap (bottom) with fiber placement. [MG, medial geniculate; 10X magnification; scale bar (100 μm)]. **(B)** Upper panel, behavioral recording paradigm; lower panel, behavioral outcome showing male and female mice spent significantly more time with the compartment containing a same-sex juvenile or opposite-sex adult, but not with the object stimulus (Two-way mixed ANOVA with Šidák’s adjusted *p=.01, ***p<.001; n = 6-8 males, 7-8 females). Open symbols denote male subjects and closed symbols denote female subjects. **(C)** Representative traces of Z scored ΔF/F of GCaMP during interactions with a same-sex juvenile (top, blue), opposite-sex adult (center, pink) and object toy mouse (bottom, yellow). **(D)** Fiber Average standardized traces from male and female mice aligned to the onset of investigation (5 sec before to 5 sec during) of a same-sex juvenile (blue), opposite-sex adult (pink) or object (yellow). **(E)** The area under the curve (AUC), calculated from average standardized traces, significantly increased during investigation of same-sex juvenile and opposite-sex adult stimuli, but not object (Two-way mixed ANOVA with Šidák’s adjusted **p<.01, ***p<.001; no sex effect; n = 6-8 males, 7-8 females; open symbols denote male subjects and closed symbols denote female subjects).

At the behavioral level, as expected, male and female mice exhibited social preference such that they spent more time investigating the compartment containing a same-sex juvenile (*M =* 203.1, *SD* = 91.33) versus empty (*M* = 100.5, *SD =* 63.72; *F*(1,13) = 9.07, *p* = .01; Fig. 2B, lower panel blue), as well as the compartment containing an opposite-sex adult (*M* = 273.3, *SD* = 81.69) versus empty (*M* = 103.8, *SD* = 61.15; *F*(1,13) = 33.14, *p <* .0001; Fig. 2B lower panel pink). For the object task, females spent more time overall investigating both the compartment containing an object and empty compartment than males (*p <* .001). However, there was no significant difference in investigation time between the compartment containing a moving toy mouse versus empty (*p* = .341; Fig. 2B, lower panel yellow).

At the neuronal activity level, we extracted and analyzed the GcaMP6m signals during each interaction with the novel stimulus (Fig. 2C). We then focused on the average neuronal activity 5 sec just before and 5 sec just after the initiation of interaction. We observed that GcaMP6m signals increased during interaction with novel same-sex or novel opposite sex stimuli, but not during interaction with a novel object (Fig. 2D). To quantify those differences, we calculated the average area under the curve for the average of each of these signals and compared between the 5 sec before and 5 sec during social interaction. The average area under the curve significantly increased in response to the same-sex juvenile from before (*M =* -0.06, *SD* = 1.26) to during investigation (*M* = 1.51, *SD* = 2.11; *F*(1, 13) = 10.68, *p* = .006; Fig. 2E blue). Similarly, activity in the PIL increased in response to the opposite-sex adult from before (*M =* 0.19, *SD* = 1.46) to during investigation (*M* = 1.78, *SD* = 1.54; *F*(1, 13) = 24.08, *p* < .001; Fig. 2E pink). However, no changes in neural activity were observed in response to investigation of the object toy mouse (*p* = .747; Fig. 2E yellow). These results demonstrate that glutamatergic neurons in the PIL specifically respond to novel social stimuli, but not a novel object, in both male and female mice.

### Glutamatergic PIL neural activity is positively associated with length of social investigation bouts, and negatively associated with the chronological order of social investigation bouts

Following our observation that the activity of glutamatergic PIL neurons was heightened in response to social stimuli (same-sex juveniles and opposite-sex adults), we performed additional analyses with a linear mixed model to understand how neural activity is associated with the length and order of social investigation bouts. We observed a positive correlation between investigation bout length and the maximum Z(dF/F) value in response to both the same-sex juvenile (beta = 0.032, *t*(221.3) = 3.93, *p* < .001; **Fig. 3A-B**) and the opposite-sex adult (beta = 0.019, *t*(200.3) = 3.88, *p* < .001; **Fig. 3D-E**). Alternatively, we observed a negative correlation between the chronological order of social investigation bouts and the maximum Z(dF/F) value in response to both the same-sex juvenile (beta = -0.069, *t*(220.2) = -3.59, *p* < .001; **Fig. 3C**) and the opposite-sex adult (beta = -0.066, *t*(203.4) = -3.70, *p* < .001; **Fig. 3F**). Together, this demonstrates that the peak of PIL neural activity increases with longer bouts of social investigation and decreases over subsequent bouts of investigation throughout the testing session.

**Figure 3.**
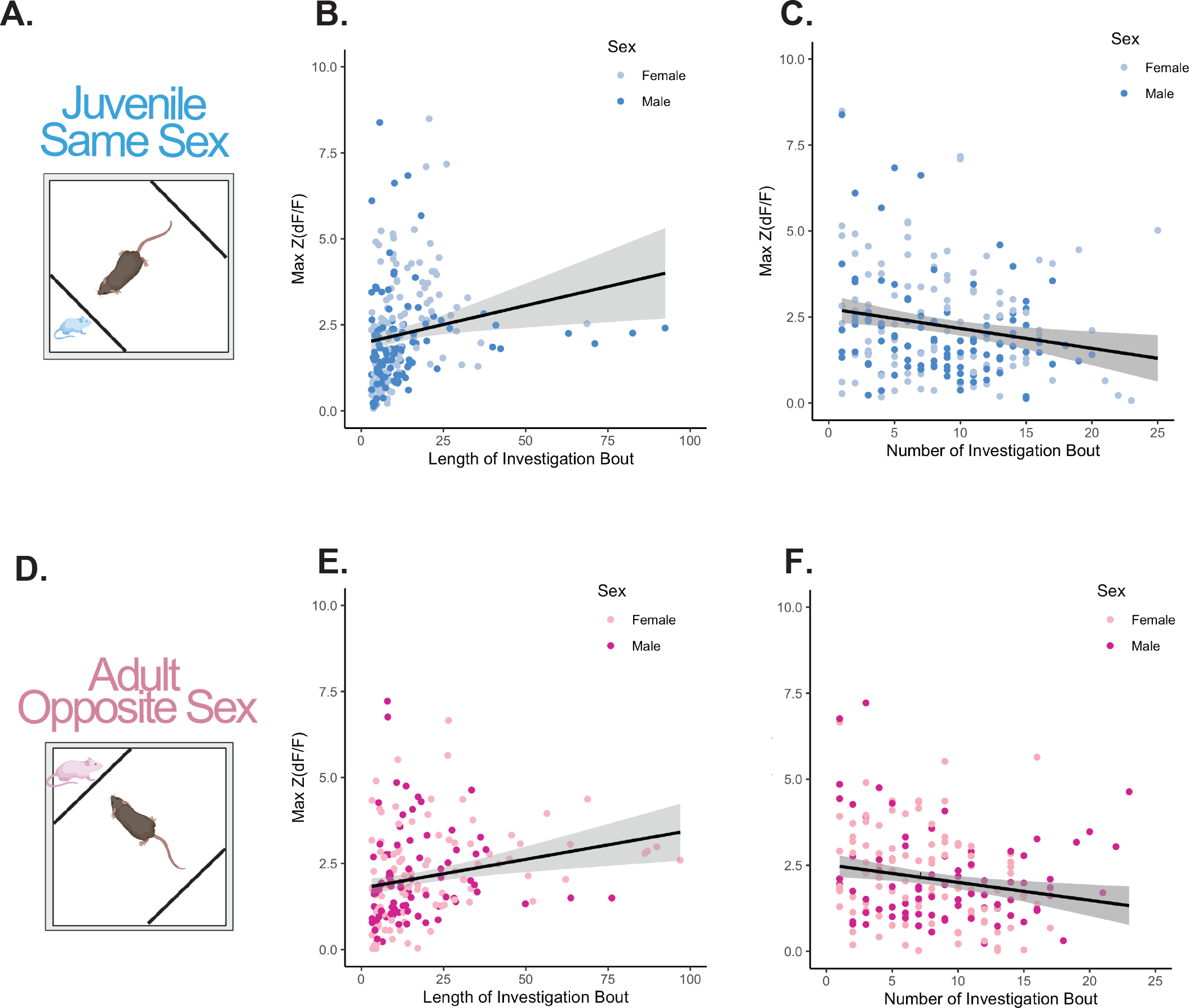
Correlations between maximum standardized signal Z(dF/F) and length of investigation bouts or chronological order of investigation bouts. **(A)** Schematic of the behavioral paradigm to record from glutamatergic neurons in the PIL during same-sex juvenile social interaction. **(B)** Positive correlation between length of investigation bout and maximum standardized signal [Max Z(dF/F)] for same-sex juvenile social interaction (linear mixed model with individual as a random factor, beta = 0.032, t(221.3) = 3.93, \*\*\**p*< .001). **(C)** Negative correlation between the chronological order of bouts and Max Z(dF/F) for same-sex juvenile social interaction (linear mixed model with individual as a random factor, beta = -0.069, t(220.2) = -3.59, \*\*\**p*< .001). **(D)** Schematic of the behavioral paradigm to record from glutamatergic neurons in the PIL during opposite-sex adult social interaction. **(E)** Positive correlation between length of investigation bout and maximum standardized signal Max Z(dF/F) for opposite-sex adult social interaction (linear mixed model with individual as a random factor, beta = 0.019, t(200.3) = 3.88, \*\*\**p*< .001). **(F)** Negative correlation between the order number of bout and Max Z(dF/F) for opposite-sex adult social interaction (linear mixed model with individual as a random factor, beta = -0.066, t(203.4) = -3.70, \*\*\**p*< .001). All data points represent individual bouts of social investigation, n = 6-7 males, 7-8 females.

### Chemogenetic inactivation of glutamatergic PIL neurons does not affect social preference in males and females

Following our observation that glutamatergic neural activity was elevated during social but not object interaction, we sought to determine whether suppression of glutamatergic PIL neurons would affect social behavior. Previous research in male and female rats using non-specific chemogenetic inhibition of PIL neurons found no effect on social novelty preference of a same-sex adult rat (11). Here, we aimed to examine how specific inhibition of PIL glutamatergic neurons impacts social preference in mice and further expand our studies to include not only a same-sex juvenile stimulus but also an opposite-sex adult. We first confirmed in a separate cohort of mice that the dose of CNO (3.5 mg/kg)(19) does not impact locomotor behavior (*p* = 0.262; Supplementary Fig. 3). We then used a dual viral chemogenetic approach with inhibitory DREADDs (AAV8-CaMKIIα-GFP-Cre + AAV2-hSyn-DIO-hM4D(Gi)-mCherry) injected bilaterally in the PIL (Fig. 4A-B). Following IP injection with CNO or vehicle, male and female mice underwent a social preference task with a novel same-sex juvenile or opposite-sex adult on separate testing days (Fig. 4C). In the juvenile social preference task, males and females preferred the same-sex juvenile to the empty chamber (*F*(1,12) = 81.25, *p* < .0001); however, there was no effect of CNO/vehicle treatment (*p* = .956; Fig. 4D-E). Additionally, there was no effect of treatment on the discrimination index (*p* = .262; Fig. 4F). Similarly, in the adult social preference task, males and females preferred the opposite-sex adult to the empty chamber (*F*(1,11) = 40.33, *p* < .0001); however, there was no effect of CNO/vehicle treatment (*p* = .096; Fig. 4G-H). Additionally, there was no effect of treatment on the discrimination index (*p* = .16; Fig. 4I). We also observed no effect of treatment with CNO on locomotor behavior in the social preference task (Supplementary Fig. 4). Together these findings suggest that although glutamatergic PIL neurons are activated during social investigation, this activity is not essential for the initiation and engagement in social interaction.

**Figure 4.**
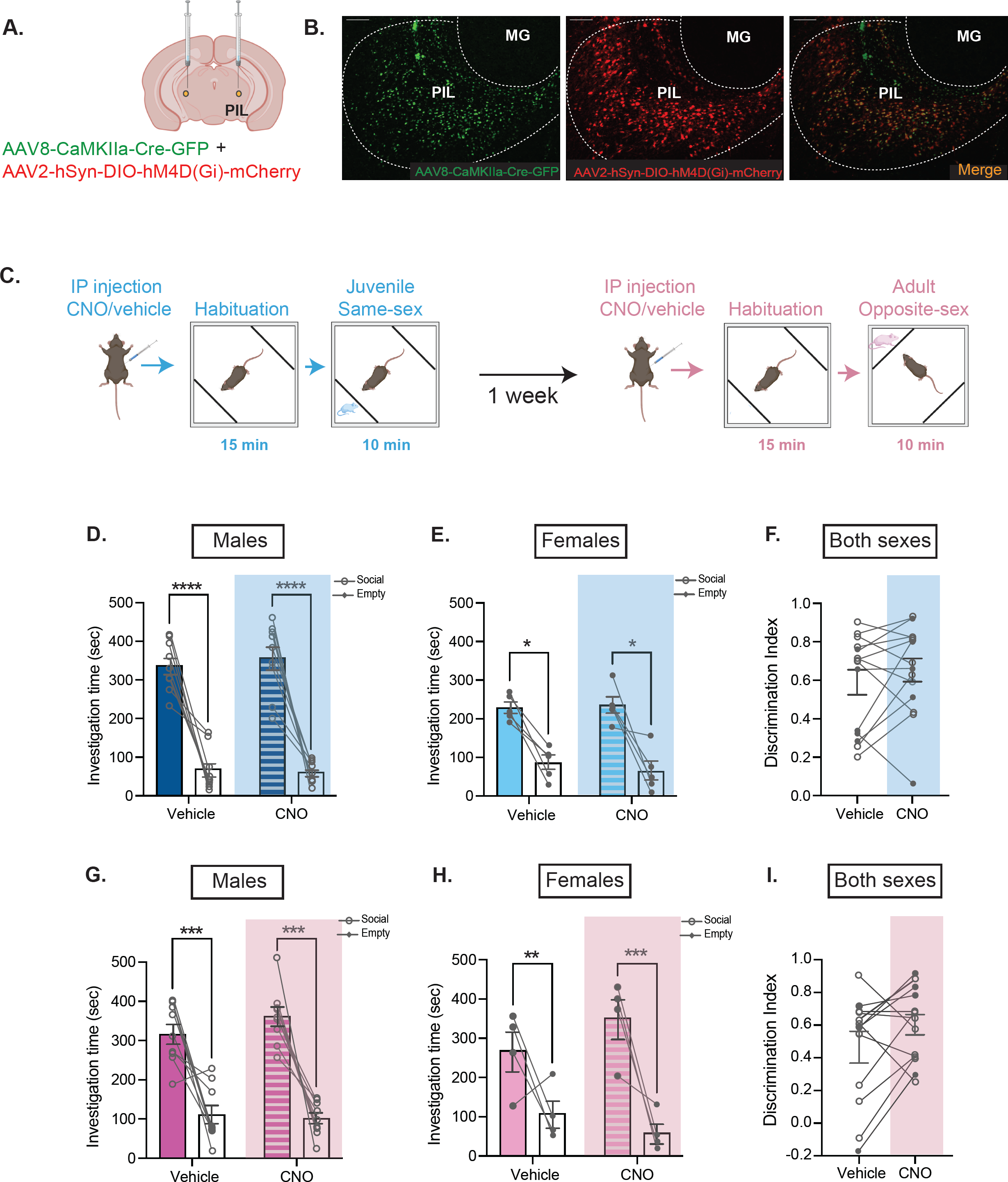
Inhibiting glutamatergic neurons in the PIL does not impact juvenile or adult social preference. **(A)** Schematic of viral injection of inhibitory DREADDs in the PIL (bilateral). **(B)** Representative histology showing CaMKIIα-GFP (left, green), hSYN-DIO-hM4D(Gi)-mCherry (middle, red) and overlap (right) with specificity to the PIL [MG, medial geniculate; 10X magnification; scale bar (100 μm)]. **(C)** Social preference behavioral paradigm following CNO/vehicle injection. **(D-E)** Males and females exhibited preference for the compartment containing the same-sex juvenile stimulus vs the empty, with no effect of CNO treatment (Three-way mixed ANOVA with Šidák’s adjusted *p<.05, ****p<.0001; n = 9 males, 5 females). **(F)** No differences were observed between treatment groups in the discrimination index ([Social – Empty]/[Social + Empty]); Two-way mixed ANOVA *p* > .05; no sex differences; n = 9 males, 5 females; open symbols denote male subjects and closed symbols denote female subjects) **(G-H)** Males and females exhibited preference for the compartment containing the opposite-sex adult social stimulus vs the empty, with no effect of CNO treatment (Three-way mixed ANOVA with Šidák’s adjusted **p<.01, ***p<.001, n = 9 males, 4 females). **(I)** No differences were observed between treatment groups in the discrimination index (Two-way mixed ANOVA *p* > .05; no sex differences; n = 9 males, 4 females; open symbols denote male subjects and closed symbols denote female subjects.).

### Chemogenetic inactivation of glutamatergic PIL neurons delays the formation of social recognition in females

Following up on our findings demonstrating increased activity of PIL glutamatergic neurons during social interaction, with decreasing peak activity over successive bouts of investigation, we hypothesized that these neurons may play a role in a different aspect of social behavior. We therefore chose to test the performance of mice following inhibition of PIL glutamatergic neurons on the social habituation-dishabituation task (21–24), which allowed us to capture two important components of social behavior: 1) engagement in free social interaction and 2) the perceptual recognition of a social stimulus over time (Fig. 5A). On this task, mice are exposed to a novel social stimulus on a first session (T1) and then exposed again to the same stimulus on three additional sessions (T2-T4). During these four sessions, the investigation time is expected to decrease, reflecting recognition of the social stimulus. In a fifth session (T5, novel), mice are introduced to a novel stimulus, to ensure that the decrease in investigation time across the previous sessions is reflecting social recognition and not lack of motivation to social interaction. Accordingly, mice are expected to spend significantly more time investigating the novel stimuli during this last session. Using this task in male mice, we found no significant changes in investigation ratio over time (T1-T4, *p* = .067) or between treatments (T1-T4, *p* = .607) (Fig. 5B) or any significant difference between treatments in investigation ratio on T5 (novel mouse) compared to T1 (*p* = .575; Fig. 5B). Furthermore, we found no effect of CNO/vehicle treatment on the total investigation time on T1 (*p* = .066; Fig. 5C) or on T5 (*p* = .684; Fig. 5D). Since the data indicate that males did not show evidence of habituation on T1-T4, even when receiving a vehicle injection, we were unable to conclude any effects of CNO treatment on performance in the social habituation-dishabituation task. However, in female mice, there was an effect of time over T1-T4 (*F*(3,24) = 3.31, *p* = .037) with a significant interaction between time and treatment (*F*(3,24) = 3.17, *p* = .043) such that female mice displayed a higher investigation ratio to the first interaction at T2 under CNO (*M* = 1.22, *SD* = 0.61) versus vehicle (*M* = 0.83, *SD* = 0.35; *t*(24) = 2.91, *p* = 0.03) and at T3 under CNO (*M* = 1.25, *SD* = 0.64) versus vehicle (*M* = 0.73, *SD* = 0.28; *t*(24) = 3.92, *p* = 0.003) but not at T4 (*p* = .799; Fig. 5E). No differences were observed in investigation ratio on T5 to T1 between treatments (*p* = .734; Fig. 5E). Additionally, there was no difference in investigation time between CNO/vehicle treatment groups in response to the novel mouse at T1 (*p* = .876; Fig. 5F) or the final novel mouse (*p* = .694; Fig. 5G). In summary, our findings demonstrate that chemogenetic inactivation of glutamatergic PIL neurons following CNO blunts the formation of social recognition in female mice.

**Figure 5.**
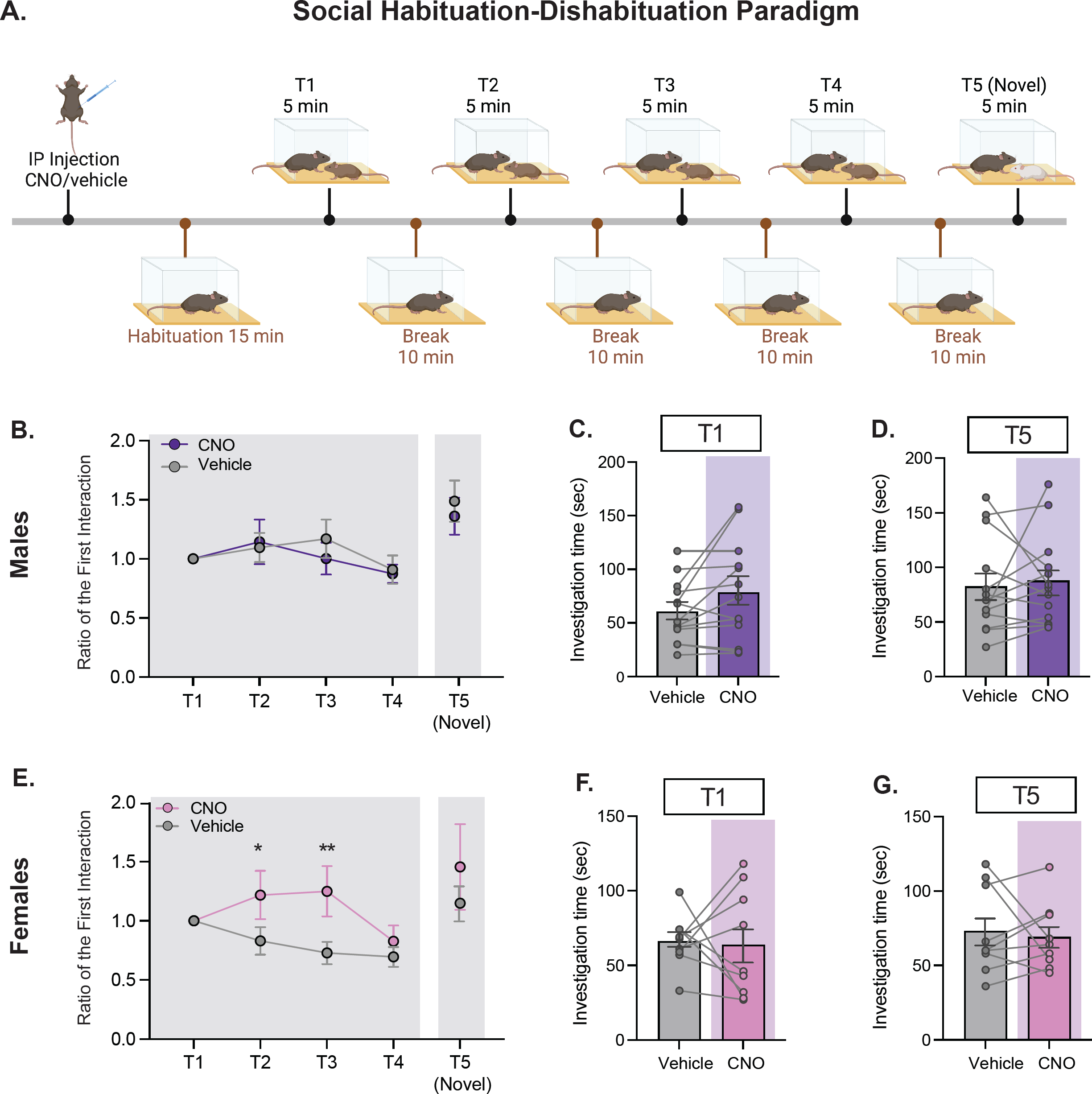
Inhibiting glutamatergic neurons in the PIL delays social recognition in females but not in males. **(A)** Social habituation-dishabituation behavioral paradigm following CNO/vehicle injection in male and female subjects injected with CaMKIIα-GFP/hSYN-DIO-hM4D(Gi)-mCherry in the PIL. **(B)** In males, no differences were observed over time or between treatment groups in ratio to the first interaction (Two-way repeated measures ANOVA *p* >.05, n=13). No differences were observed between contact ratio of the novel mouse to T1 (Paired samples t-test *p*>.05, n = 13) **(C)** No differences were observed in males in investigation time between treatment groups at T1 (Paired samples t-test *p* >.05, n=13). **(D)** No differences were observed in males in investigation time between treatment groups at the presentation of the final novel mouse (Paired samples t-test *p* >.05, n=13). **(E)** In females, there was a significant interaction between treatment and time (Two-way repeated measures ANOVA \**p*<.05) such that female mice displayed a higher contact ratio to the first interaction under CNO at T2 (Šidák’s adjusted \**p*<.05) and at T3 (Šidák’s adjusted \*\**p* < .01) but not T4 (Šidák’s adjusted *p* >.05). No differences were observed between contact ratio of the novel mouse to T1 (Wilcoxon matched-pairs signed rank test *p*>.05, n = 9). **(F)** No differences were observed in females in investigation time between treatment groups at T1 (Paired samples t-test *p* >.05, n=9). **(G)** No differences were observed in females in investigation time between treatment groups at the presentation of the final novel mouse (Paired samples t-test *p* >.05, n=9).

## DISCUSSION

Social interaction is a dynamic sequence of events that requires detection and efficient integration of social sensory cues (29). The PIL, located at the posterior thalamus, is well-positioned to convey sensory cues to the brain social network to modulate social behavior (1). For example, studies on the PIL in the context of maternal reflexes have shown that *c-fos*-immunoreactivity of PIL neurons, particularly TIP39^+^ neurons, is increased in maternal rats following suckling (2, 4). Based on neuroanatomical studies in lactating rats showing that TIP39^+^ neurons project to the MPOA (4, 30) and the PVH (3), it has been suggested that sensory information from pups during maternal reflexes and behavior is conveyed to the MPOA and PVH of the lactating mothers through the PIL➔MPOA and/or PIL➔PVH pathways. It has also been demonstrated that the PIL➔MPOA pathway is involved in grooming between conspecific female rats during social interaction, therefore playing a role in affiliative social interaction (11). A recent study conducted in mice focused on the subparafascicular nucleus (SPFp), which shares neuroanatomical borders with the PIL, and identified a population of SPFp neurons that express calcitonin gene-related peptide (CGRP). This study revealed that CGRP expressing neurons respond to multi-sensory threat stimuli and project to the lateral amygdala to regulate fear memory (26). Together, the results from rat and mouse studies suggest that the PIL plays a role in processing sensory inputs, although the specific neural pathways involved may differ depending on the species, sex, and behavioral context.

Here we show that in both male and female mice, PIL glutamatergic neurons are also activated in response to social interactions with a novel social stimulus and that their response decreases over time as social stimuli become less novel, which could point to a role in conveying the salience of a social stimulus. Furthermore, we demonstrate that this activation occurs during both same-sex and opposite-sex social interactions, suggesting that the PIL may function to communicate information about social stimuli not only during maternal and same-sex behaviors but also during opposite sex behavior. The role of the PIL in sexual behavior is supported by earlier work showing increased *c-fos*-immunoreactivity following ejaculation in male rats and vaginocervical stimulation in female rats in the SPFp (31–33).

Using targeted chemogenetic tools, we show that inhibiting glutamatergic PIL neurons does not impact the mice’s ability to form social preference of a same-sex juvenile stimulus. This observation is in line with recent findings from Keller and colleagues in male and female rats, which showed that rats maintained social preference for a novel conspecific versus familiar cage mate following inhibition of PIL neurons (11). These findings suggest that the increase in activity of PIL glutamatergic neurons, which we observed during constrained social interaction, is not critical for the initiation and/or engagement of social interaction, but may rather play a role in conveying sensory information about the identity and/or the salience of the social stimulus, thus facilitating social recognition memory (1), which is the ability of the animal to recognize other conspecifics (34).

To address this proposition, we chose the habituation dishabituation task (21–24), which allowed us to examine the effect of PIL inhibition on social interaction and/or social recognition within the same paradigm. While our data in males were not conclusive, possibly due to increased aggressive behavior, in females, we found that initial social interaction with a novel stimulus was not impacted by the inhibition of PIL glutamatergic neurons (no difference between CNO and vehicle treatment on T1 or T5), thus confirming that PIL activity is not required for social interaction per-se. However, we found that inhibition of glutamatergic PIL neurons delayed the time to habituation (CNO treatment led to a higher ratio of investigation at T2 and T3), suggesting a delay in social recognition. Notably, our observation that social recognition is eventually achieved (on T4), indicates that glutamatergic neurons of the PIL are not the only player in encoding social sensory and/or somatosensory information during social recognition and that other neuronal populations of the PIL and/or brain regions may be involved and can eventually compensate for the lack of PIL glutamatergic activity, yet with some delay. For instance, structures such as the lateral septum, hippocampus, amygdala, and medial prefrontal cortex have been identified as key players in the social recognition memory network (35–38). These regions may receive sensory information via other brain regions and operate within pathways separate from the PIL that could compensate for the decrease in neural activity of PIL glutamatergic neurons, yet with a delay. Furthermore, considering the fact that PIL neurons project to OXT neurons of the PVH and OXT is known to be involved in social recognition memory (13, 21, 34; 39-44), it is possible that inhibition of neural activity of PIL glutamatergic neurons could affect social recognition memory by reducing the release of OXT. This, in turn, could potentially hinder the processing of the social stimulus salience and delay the formation of social recognition memory, a possibility which should be explored in forthcoming research.

Together our findings suggest that glutamatergic PIL neurons are involved in communicating sensory information to promptly mediate social recognition. Future research should be aimed at dissecting the contribution of specific glutamatergic PIL pathways in social recognition and social behaviors in general, with consideration of possible sexual dimorphism in the function of the PIL in different behavioral contexts. We believe that such studies will be of translational significance for brain disorders that are characterized by social behavior and social recognition deficits, including autism spectrum disorder (45–48), where several autism-risk genes were reported to encode for glutamatergic synaptic components and to impact social recognition memory (49–51). Of relevance would be the PIL➔PVH pathway, specifically glutamatergic inputs from the PIL to the oxytocin neurons of the PVH, given the role of glutamatergic circuits within the hypothalamus in the release of oxytocin (52) and the well-established role of oxytocin in social behaviors and social recognition (13, 21, 34; 39-44).

## ACKNOWLEDGEMENT

The work reported in this publication was supported by NIMH F31MH129025 and a Seaver Foundation fellowship to A.B.L and H.H.N. We thank Dr. Eric Nestler for sharing fiber photometry equipment and resources and Mr. Nicholas Cordero for assisting with cell counting.

## DISCLOSURES

Ms. Amanda Leithead, Dr. Arthur Godino, Dr. Marie Barbier, and Dr. Harony-Nicolas report no biomedical financial interests or potential conflicts of interest.

## Notes

### Competing Interest Statement

The authors have declared no competing interest.

## REFERENCES

1. Dobolyi A, Cservenák M, Young LJ (2018): Thalamic integration of social stimuli regulating parental behavior and the oxytocin system. Frontiers in Neuroendocrinology. 51:102–115.

2. Cservenák M, Bodnár I, Usdin TB, Palkovits M, Nagy GM, Dobolyi A (2010): Tuberoinfundibular peptide of 39 residues is activated during lactation and participates in the suckling-induced prolactin release in rat. Endocrinology. 151:5830–5840.

3. Cservenák M, Keller D, Kis V, Fazekas EA, Öllös H, Lékó AH, et al. (2017): A thalamo-hypothalamic pathway that activates oxytocin neurons in social contexts in female rats. Endocrinology. 158:335–348.

4. Cservenák M, Szabó ÉR, Bodnár I, Lékó A, Palkovits M, Nagy GM, et al. (2013): Thalamic neuropeptide mediating the effects of nursing on lactation and maternal motivation. Psychoneuroendocrinology. 38:3070–3084.

5. Hansen S, Köhler C (1984): The importance of the peripeduncular nucleus in the neuroendocrine control of sexual behavior and milk ejection in the rat. Neuroendocrinology. 39:563–572.

6. Factor EM, Mayer AD, Rosenblatt JS (1993): Peripeduncular nucleus lesions in the rat: I. Effects on maternal aggression, lactation, and maternal behavior during pre-and postpartum periods. Behavioral Neuroscience. 107:166.

7. Cai D, Yue Y, Su X, Liu M, Wang Y, You L, et al. (2019): Distinct anatomical connectivity patterns differentiate subdivisions of the nonlemniscal auditory thalamus in mice. Cerebral Cortex. 29:2437–2454.

8. Ledoux JE, Ruggiero DA, Forest R, Stornetta R, Reis DJ (1987): Topographic organization of convergent projections to the thalamus from the inferior colliculus and spinal cord in the rat. Journal of Comparative Neurology. 264:123–146.

9. Linke R (1999): Differential projection patterns of superior and inferior collicular neurons onto posterior paralaminar nuclei of the thalamus surrounding the medial geniculate body in the rat. European Journal of Neuroscience. 11:187–203.

10. Yasui Y, Kayahara T, Nakano K, Mizuno N (1990): The subparafascicular thalamic nucleus of the rat receives projection fibers from the inferior colliculus and auditory cortex. Brain research. 537:323–327.

11. Keller D, Láng T, Cservenák M, Puska G, Barna J, Csillag V, et al. (2022): A thalamo-preoptic pathway promotes social grooming in rodents. Current Biology. 32: 4593–4606.

12. Tang Y, Benusiglio D, Lefevre A, Hilfiger L, Althammer F, Bludau A, et al. (2020): Social touch promotes interfemale communication via activation of parvocellular oxytocin neurons. Nature Neuroscience. 23:1125–1137.

13. Bielsky IF, Young LJ (2004): Oxytocin, vasopressin, and social recognition in mammals. Peptides. 25:1565–1574.

14. Donaldson ZR, Young LJ (2008): Oxytocin, vasopressin, and the neurogenetics of sociality. Science. 322:900–904.

15. Ferguson JN, Young LJ, Insel TR (2002): The neuroendocrine basis of social recognition. Frontiers in Neuroendocrinology. 23:200–224.

16. Insel TR, Young LJ (2001): The neurobiology of attachment. Nature Reviews Neuroscience. 2:129–136.

17. Insel TR, Young LJ (2000): Neuropeptides and the evolution of social behavior. Current Opinion in Neurobiology. 10:784–789.

18. Froemke RC, Young LJ (2021): Oxytocin, neural plasticity, and social behavior. Annual Review of Neuroscience. 44:359–381.

19. Jendryka M, Palchaudhuri M, Ursu D, van der Veen B, Liss B, Kätzel D, et al. (2019): Pharmacokinetic and pharmacodynamic actions of clozapine-N-oxide, clozapine, and compound 21 in DREADD-based chemogenetics in mice. Scientific Reports. 9:1–14.

20. Netser S, Haskal S, Magalnik H, Wagner S (2017): A novel system for tracking social preference dynamics in mice reveals sex- and strain-specific characteristics. Molecular Autism. 8:53.

21. Ferguson JN, Young LJ, Hearn EF, Matzuk MM, Insel TR, Winslow JT (2000): Social amnesia in mice lacking the oxytocin gene. Nature Genetics. 25:284–288.

22. Cao Y, Wu R, Tai F, Zhang X, Yu P, An X, et al. (2014): Neonatal paternal deprivation impairs social recognition and alters levels of oxytocin and estrogen receptor α mRNA expression in the MeA and NAcc, and serum oxytocin in mandarin voles. Hormones and Behavior. 65:57–65.

23. Spiteri T, Ågmo A (2009): Ovarian hormones modulate social recognition in female rats. Physiology & Behavior. 98:247–250.

24. Jacobs SA, Huang F, Tsien JZ, Wei W (2016): Social recognition memory test in rodents. Bio-protocol. 6:e1804–e1804.

25. Morel C, Montgomery SE, Li L, Durand-de Cuttoli R, Teichman EM, Juarez B, et al. (2022): Midbrain projection to the basolateral amygdala encodes anxiety-like but not depression-like behaviors. Nature Communications. 13:1–13.

26. Kang SJ, Liu S, Ye M, Kim D-I, Pao GM, Copits BA, et al. (2022): A central alarm system that gates multi-sensory innate threat cues to the amygdala. Cell Reports. 40:111222.

27. Norman KJ, Riceberg JS, Koike H, Bateh J, McCraney SE, Caro K, et al. (2021): Post-error recruitment of frontal sensory cortical projections promotes attention in mice. Neuron. 109:1202–1213. e1205.

28. Muir J, Lorsch ZS, Ramakrishnan C, Deisseroth K, Nestler EJ, Calipari ES, et al. (2018): In vivo fiber photometry reveals signature of future stress susceptibility in nucleus accumbens. Neuropsychopharmacology. 43:255–263.

29. Chen P, Hong W (2018): Neural Circuit Mechanisms of Social Behavior. Neuron. 98:16–30.

30. Szabo FK, Snyder N, Usdin TB, Hoffman GE (2010): A direct neuronal connection between the subparafascicular and ventrolateral arcuate nuclei in non-lactating female rats. Could this pathway play a role in the suckling-induced prolactin release? Endocrine. 37:62–70.

31. Coolen LM, Peters HJ, Veening JG (1996): Fos immunoreactivity in the rat brain following consummatory elements of sexual behavior: a sex comparison. Brain Research. 738:67–82.

32. Coolen L, Peters H, Veening J (1997): Distribution of Fos immunoreactivity following mating versus anogenital investigation in the male rat brain. Neuroscience. 77:1151–1161.

33. Veening JG, Coolen LM (1998): Neural activation following sexual behavior in the male and female rat brain. Behavioural Brain Research. 92:181–193.

34. Insel TR, Fernald RD (2004): How The Brain Processes Social Information: Searching for the Social Brain. Annual Review of Neuroscience. 27:697–722.

35. Ferguson JN, Aldag JM, Insel TR, Young LJ (2001): Oxytocin in the medial amygdala is essential for social recognition in the mouse. Journal of Neuroscience. 21:8278–8285.

36. Gur R, Tendler A, Wagner S (2014): Long-Term Social Recognition Memory Is Mediated by Oxytocin-Dependent Synaptic Plasticity in the Medial Amygdala. Biolical Psychiatry.

37. Tanimizu T, Kenney JW, Okano E, Kadoma K, Frankland PW, Kida S (2017): Functional connectivity of multiple brain regions required for the consolidation of social recognition memory. Journal of Neuroscience. 37:4103–4116.

38. Wang X, Zhan Y (2022): Regulation of social recognition memory in the hippocampal circuits. Frontiers in Neural Circuits. 16.

39. Lin YT, Hsieh TY, Tsai TC, Chen CC, Huang CC, Hsu KS (2018): Conditional deletion of hippocampal CA2/CA3a oxytocin receptors impairs the persistence of long-term social recognition memory in mice. Journal of Neuroscience. 38:1218–1231.

40. Lukas M, Toth I, Veenema AH, Neumann ID (2013): Oxytocin mediates rodent social memory within the lateral septum and the medial amygdala depending on the relevance of the social stimulus: male juvenile versus female adult conspecifics. Psychoneuroendocrinology. 38:916–926.

41. Penn DJ, Frommen JG (2010): Kin recognition: an overview of conceptual issues, mechanisms and evolutionary theory. Animal Behaviour: Evolution and Mechanisms. 55–85.

42. Popik P, Vetulani J, Van Ree JM (1992): Low doses of oxytocin facilitate social recognition in rats. Psychopharmacology. 106:71–74.

43. Raam T, McAvoy KM, Besnard A, Veenema AH, Sahay A (2017): Hippocampal oxytocin receptors are necessary for discrimination of social stimuli. Nature communications. 8:1–14.

44. Takayanagi Y, Yoshida M, Bielsky IF, Ross HE, Kawamata M, Onaka T, et al. (2005): Pervasive social deficits, but normal parturition, in oxytocin receptor-deficient mice. Proceedings of the National Academy of Sciences. 102:16096–16101.

45. American Psychiatric Association D (2013): American Psychiatric Association Diagnostic and Statistical Manual of Mental Disorders.

46. Williams DL, Goldstein G, Minshew NJ (2005): Impaired memory for faces and social scenes in autism: Clinical implications of memory dysfunction. Archives of Clinical Neuropsychology. 20:1–15.

47. Minio-Paluello I, Porciello G, Pascual-Leone A, Baron-Cohen S (2020): Face individual identity recognition: a potential endophenotype in autism. Molecular Autism. 11:1–16.

48. Weigelt S, Koldewyn K, Kanwisher N (2012): Face identity recognition in autism spectrum disorders: A review of behavioral studies. Neuroscience & Biobehavioral Reviews. 36:1060–1084.

49. De Rubeis S, He X, Goldberg AP, Poultney CS, Samocha K, Cicek AE, et al. (2014): Synaptic, transcriptional and chromatin genes disrupted in autism. Nature. 515:209–215.

50. Satterstrom FK, Kosmicki JA, Wang J, Breen MS, De Rubeis S, An J-Y, et al. (2020): Large-scale exome sequencing study implicates both developmental and functional changes in the neurobiology of autism. Cell. 180:568–584. e523.

51. Harony-Nicolas H, Kay M, Hoffmann JD, Klein ME, Bozdagi-Gunal O, Riad M, et al. (2017): Oxytocin improves behavioral and electrophysiological deficits in a novel Shank3-deficient rat. Elife. 6:e18904.

52. Leithead AB, Tasker JG, Harony-Nicolas H (2021): The interplay between glutamatergic circuits and oxytocin neurons in the hypothalamus and its relevance to neurodevelopmental disorders. Journal of Neuroendocrinology. 33:e13061.

