## Supplement for "Social Interaction Elicits Activity in Glutamatergic Neurons in the Posterior Intralaminar Complex of the Thalamus"

**Supplemental Methods and Materials:**

Stereotaxic surgery

Mice were anesthetized with 5% isoflurane for induction and then maintained at 1.5–2.5% isoflurane with 2% oxygen using a tabletop vaporizer and a non-breathing circuit. Aseptic surgical techniques were followed. The virus was injected into the PIL (A–P -3.1 mm, M–L +/- 1.8 mm, D–V -3.6 mm) or PVH (A-P -0.6 mm, M-L +/- 1 mm, D–V -5.15 mm, 10-degree angle) with a 10 µl 33G NanoFil syringe (World Precision Instruments, Sarasota, FL, USA) at a rate of 0.1 µl/min for 0.3 µl total volume. The syringe was then left in place for an additional 10 min and afterward withdrawn at a rate of 0.2 mm/min. For fiber photometry experiments, a fiber optic cannula (400 µm 0.39NA, Cat. #CFM14L05-10, Thorlabs, Newton, NJ, USA) was implanted into the PIL, using the same coordinates as for the viral injection, and secured to the skull with dental cement (Cat. #51459, Stoelting Co., Wood Dale, IL, USA). Incision wounds were closed using sutures (Ethilon Suture 5-0, Henry Schein, Melville, NY, USA). Mice received intraoperative subcutaneous fluids for hydration (Lactated Ringer Solution, Thermo Fisher Scientific, Waltham, MA, USA) and buprenorphine (0.05 mg/kg) was administered every 12 hours for 72 hours post-operatively.

Histology

Mice were anesthetized with 5% isoflurane, then perfused with 0.2 M Sodium Phosphate Buffer followed by 4% PFA at a rate of 8 mL/min for 10 min. Brains were removed and placed in 4% PFA overnight at 4 °C, then immersed in a sucrose solution (30% sucrose, 100 mM glycine and 0.05% sodium azide in 1×PBS) for 48 hours at 4 °C. Brains were frozen in a cryomold filled with

O.C.T. (Tissue-Tek, Torrance, CA) and stored at – 80 °C until sectioning with a cryostat (Leica CM 1860 Leica Biosytems, Buffalo Grove, IL, USA).

*c-fos* immunohistochemistry

Immunohistochemistry for *c-fos* was performed on 40 μm sections containing the PIL (bregma – 2.8 to 3.4 mm A-P). 13-15 sections per mouse were collected and stained for *c-fos*. Sections were washed (3 × 10 min each in 1×PBS + 0.05% Triton X-100) then blocked and permeabilized for 1 hour in 5% donkey serum (Jackson ImmunoResearch, West Grove, PA, USA) in 1XPBS + 0.5% Triton X-100. Anti-*c-fos* (9F6) rabbit monoclonal antibody (Cat. #2250, Cell Signaling, Danvers, MA, USA) was then applied to sections in blocking solution at a concentration of 1:3,000 and left to incubate for 48 hours at 4 °C. Sections were washed then incubated with 1:1,000 donkey anti-rabbit 594 (Cat. # A21207, Invitrogen, Waltham, MA, USA) for 2 hours at room temperature, after which they were washed again and mounted with DAPI (Cat. #H-1200, Vector Labs, Burlingame, CA, USA) for microscopy and cell counting.

RNAscope

We performed two separate RNAscope Multiplex Fluorescent assays on PIL brain sections: 1) mice injected with AAV1-hSyn-eGFP-Cre in the PVH to determine whether projections from the PVH to the PIL were glutamatergic/GABAergic, and 2) mice injected with AAV8-CaMKIIα-GFP in the PIL to determine whether PIL neurons expressing CaMKIIα are glutamatergic/GABAergic. Mouse GFP (Cat. #400281), *vglut2* (*slc17a6;* Cat. #319171) and *vgat1* (*slc32a1;* Cat. #319191) probes were used (ACDBio, Newark, CA, USA). Mice were deeply anesthetized with 5% isoflurane then underwent cervical decapitation and brain collection. Fresh brains were flash frozen in isopentane over dry ice, then immediately sectioned at 15 μm, mounted onto glass slides (SuperFrost Plus Microscope Slides, Fisher Scientific) and stored at -80°C until the start of the RNAscope procedure. The manufacturer's protocol (RNAscope Multiplex Fluorescent Reagent Kit, ACDBio) was followed: Tissue sections were first thawed at room temperature (RT) for 10 minutes, then fixed with 4% PFA for 15 minutes at 4°C, and then dehydrated with ethanol at RT. Sections were then incubated in hydrogen peroxide ($H_2O_2$) for 10 minutes. The GFP, *vglut2* and *vgat1* probes were mixed with probe dilutant (1:1:1:50) and added to the sections, then left to incubate for 2 hours in a 40°C oven (HybEZ II Hybridization System, ACDBio). This was followed by amplification with probes (Amp1, Amp2, and Amp3), provided by the kit. Finally, sections were incubated with opal dyes (Akoya Biosciences, Marlborough, MA, USA) 520, 570, and 690 and cover slipped with VECTASHIELD Antifade Mounting Medium with DAPI.

<u>Behavioral Assays</u>

*Social/Object Preference*

To measure preference, mice were habituated to an open field box with two empty compartments on opposite corners of the box for 15 minutes. In the social preference task (1) a social stimulus (same-sex juvenile or opposite-sex adult) was then placed into one compartment with the opposite compartment left empty (counterbalanced). Compartments had a wire mesh window which allowed for the exchange of visual, auditory, and olfactory information, but restricted physical touch. Similarly, in the object preference task a toy mouse was placed into a compartment with a string which allowed the object to rotate, to control for the effect of motion on the subject mouse's attention and motivation to interact. The subject mouse was then allowed to interact for 10 minutes.

Investigation time of the stimulus-containing chamber and empty chamber was captured and converted to a discrimination index ([Stimulus – Empty]/[Stimulus + Empty]).

*Social Habituation-Dishabituation*

In the social habituation-dishabituation paradigm (2-5), the subject mouse was first habituated to an empty open field box for 15 minutes. A novel same-sex juvenile was then placed into the box and free social interaction was permitted for 5 minutes (T1). The juvenile was then removed from the box, leaving the subject mouse alone in the box for a 10-minute break. This series was then repeated 3 more times with the same juvenile (T2-T4) to capture habituation to the stimulus. A different novel same-sex juvenile was then introduced for one final 5-minute bout (T5) to control for lack of motivation to engage in social interaction. Investigation time between stimuli was captured for each timepoint and calculated as ratios to the first interaction for each mouse (baseline social interaction per mouse). For example, of T3 = [investigation time T3]/[investigation time T1].

Fiber Photometry Recording

Behavior and calcium activity were recorded using methods previously described (6). The fiber photometry system and video-tracking system (Ethovision XT 11, Noldus, Leesburg, VA, USA) were time-locked with transistor–transistor logic signals (TTLs). The fiber photometry system used two light-emitting diodes at 490 and 405 nm (Thorlabs), reflected off dichroic mirrors (Cat. #FF495; Semrock, West Henrietta, NY, USA) and coupled into a 400-µm 0.57 N.A. fiberoptic patchcord (Cat. #MFP_400/430/1100-0.57; Doric, Quebec, QC, CA). The real-time fiber photometry signal was collected using a signal processer (Tucker–Davis Technologies) and acquired with open source OpenEx software 2.20 controlling an RX8 lock-in amplifier (Tucker-

Davis Technologies, Alachua, FL, USA). OpenEx (https://www.tdt.com/support/downloads/ ; and https://www.tdt.com/component/openex-software-suite/), sinusoidally modulated each LED's output (490 nm at 211 Hz, and 405 nm isosbestic control at 531 Hz). The two output signals were then projected onto a photodetector (2151 femtowatt photoreceiver; Newport, Irvine, CA, USA). The photoreceiver signal was sampled at 6.1 kHz, after which each of the two modulated signals was separated by the real-time processor for analysis. Decimated signals were collected at a sampling frequency of 381 Hz to perform the post-acquisition analyses.

### Analyses

#### *Fiber Photometry Analyses*

Post-acquisition analyses were performed using custom programs. First, TDT data were extracted and converted into fluorescence time-series using generic MATLAB code from the Lerner lab (https://github.com/talialerner/). To compare neuronal activity across animals and behavioral sessions, individual animal time-series data were analyzed using custom R codes following the standard method previously detailed (7), with minor modifications. Briefly, both 490 and 405 nm signals were first smoothed using a $4^{th}$-order 5Hz lowpass Butterworth filter built using the gsignal::butter function. To remove the bleaching slope and low-frequency fluctuations, baseline correction was then performed by subtracting the baseline obtained by regressing each individual signal using the LOWESS smoother (stats::lowess) with default parameters from the smoothed 490 and 405 nm signals. Both 490 and 405 signals were then standardized using a robust z-score (zF = (F – median(F))/mad(F)). The standardized 405 nm signal was then fitted to the standardized 490 nm signal using the robust regression function MASS::rlm, and normalized dF/F (z(dF/F) was finally calculated as the difference between the 490 nm signal and the fitted 405 nm signal to remove motion artifacts and autofluorescence. To analyze time-locked neuronal activity in respect to behavior, the normalized z(dF/F) signal was extracted from 5 sec to 5 sec around the onset of the relevant behavior (defined as $t = 0$ sec). Before averaging, each epoch was offset such that each z-score average from -5 to 5 equaled zero. Finally, signal changes was quantified for relevant time intervals as the corresponding areas under the curve, which were calculated with linear interpolation using the MESS::auc function. For additional analyses, the maximum peak of the normalized z(dF/F) signal for each bout of stimulus interaction was calculated using the max function.

*Behavioral Analyses*

Behavioral videos were recorded (C615 Webcam, Logitech, Lusanne, CH) and analyzed using TrackRodent, an open-source Matlab-based automated tracking system that uses a body-based algorithm (1). The source code is available on GitHub (https://github.com/shainetser/TrackRodent). For analysis of the social/object preference task behavior videos, the area around the wire mesh portion of the chamber was highlighted so that 'investigation' was calculated as the duration of time in which the subject mouse's body was in contact with the highlighted interaction zone. For analysis of the social habituation-dishabituation behavior videos, 'investigation' was calculated as the duration of time in which the subject mouse and stimulus mouse were in physical contact.

*Microscopy Analyses*

Viral expression, fiber placement, and immunohistochemical staining were visualized with an EVOS imaging system (Thermo Fisher Scientific) at 4x and 10x magnification. For *c-fos* experiments, images were exported to FIJI (ImageJ) and a boundary was drawn around the PIL. *C-fos*$^+$ cells were then manually counted by a scorer blind to experimental conditions using the cell counter feature on FIJI. The area ($mm^2$) of each image was calculated using the FIJI program to determine the number of *c-fos*$^+$ cells per $mm^2$. The area per section ranged from approximately 0.15 mm to 0.5 mm. For fluorescent RNAscope images, a Zeiss AxioImager Z2M with ApoTome.2 at 10x and 40x magnification was used at the Microscopy Core at the Icahn School of Medicine at Mount Sinai.

Statistical analyses were performed using GraphPad Prism 9.0 software and image graphics were created using BioRender.com and Adobe Illustrator.

**Supplementary Figure 1.**

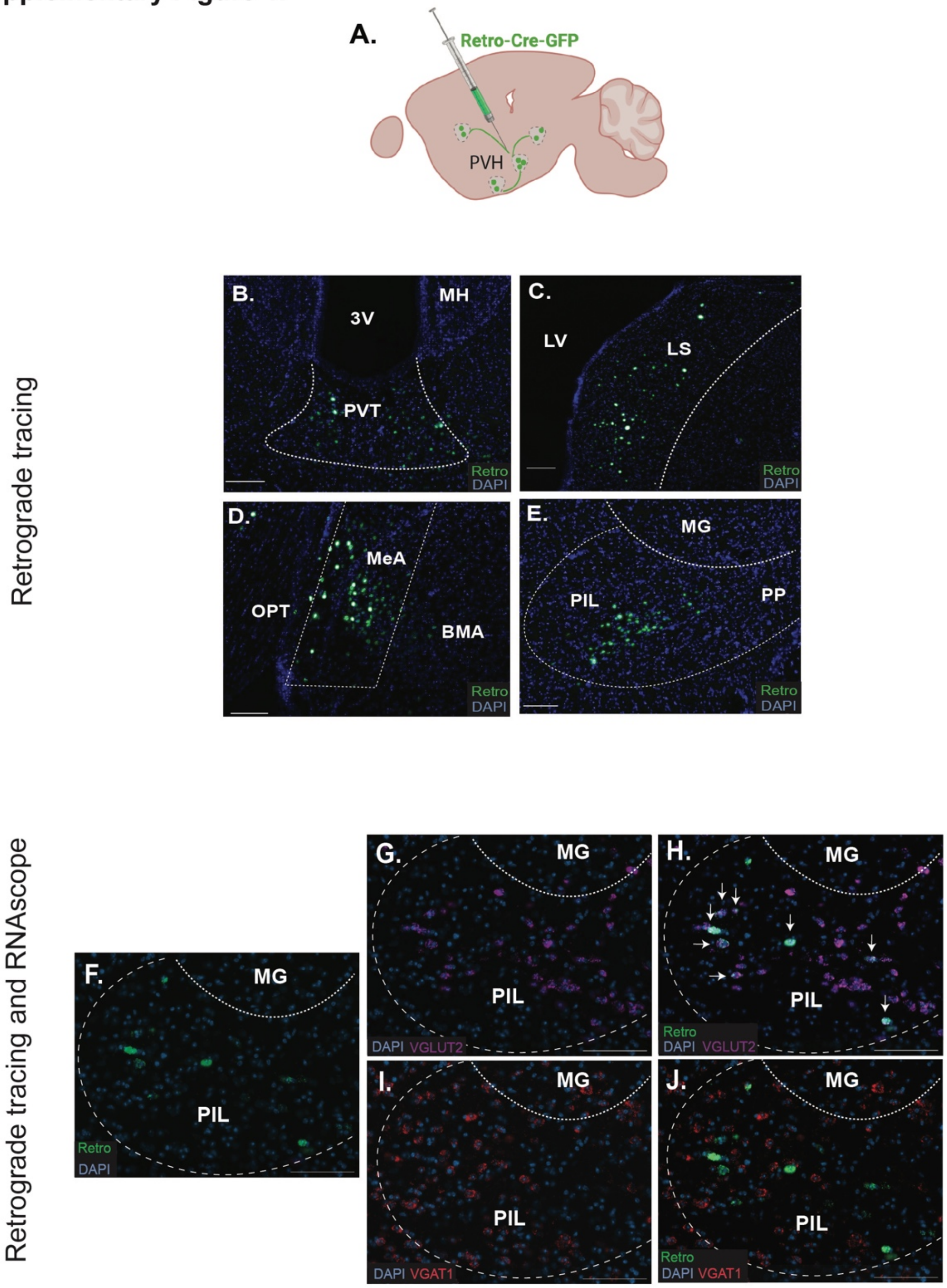

**Supplementary Figure 1. Several regions project to the PVH, including the PIL which sends excitatory, but not inhibitory, monosynaptic inputs. (A)** Schematic of monosynaptic retrograde viral tracing from the PVH of the hypothalamus with a GFP-tagged Retro-Cre virus. **(B-E)** Retrograde GFP expression was observed in the paraventricular nucleus of the thalamus (PVT), the lateral septum (LS), the medial amygdala (MeA), and the posterior intralaminar complex of the thalamus (PIL) [10X magnification; scale bar (100 μm)]. **(F-J)** RNAscope performed on PIL sections containing Retro-GFP (green) for *vglut2* (violet) revealed overlap with Retro-GFP$^{+}$ cells, indicating that some projections from the PIL to the PVH are glutamatergic. RNAscope for *vgat1* (red) revealed no overlap with Retro-GFP$^{+}$ cells, indicating a lack of GABAergic projections from the PIL to the PVH. [20X magnification; scale bar (100 μm); 3V, third ventricle; LV, lateral ventricle; OPT, optic tract; BMA, basomedial amygdala; MG, medial geniculate; SN, substantia nigra; PVH, paraventricular nucleus of the hypothalamus].

**Supplementary Figure 2.**

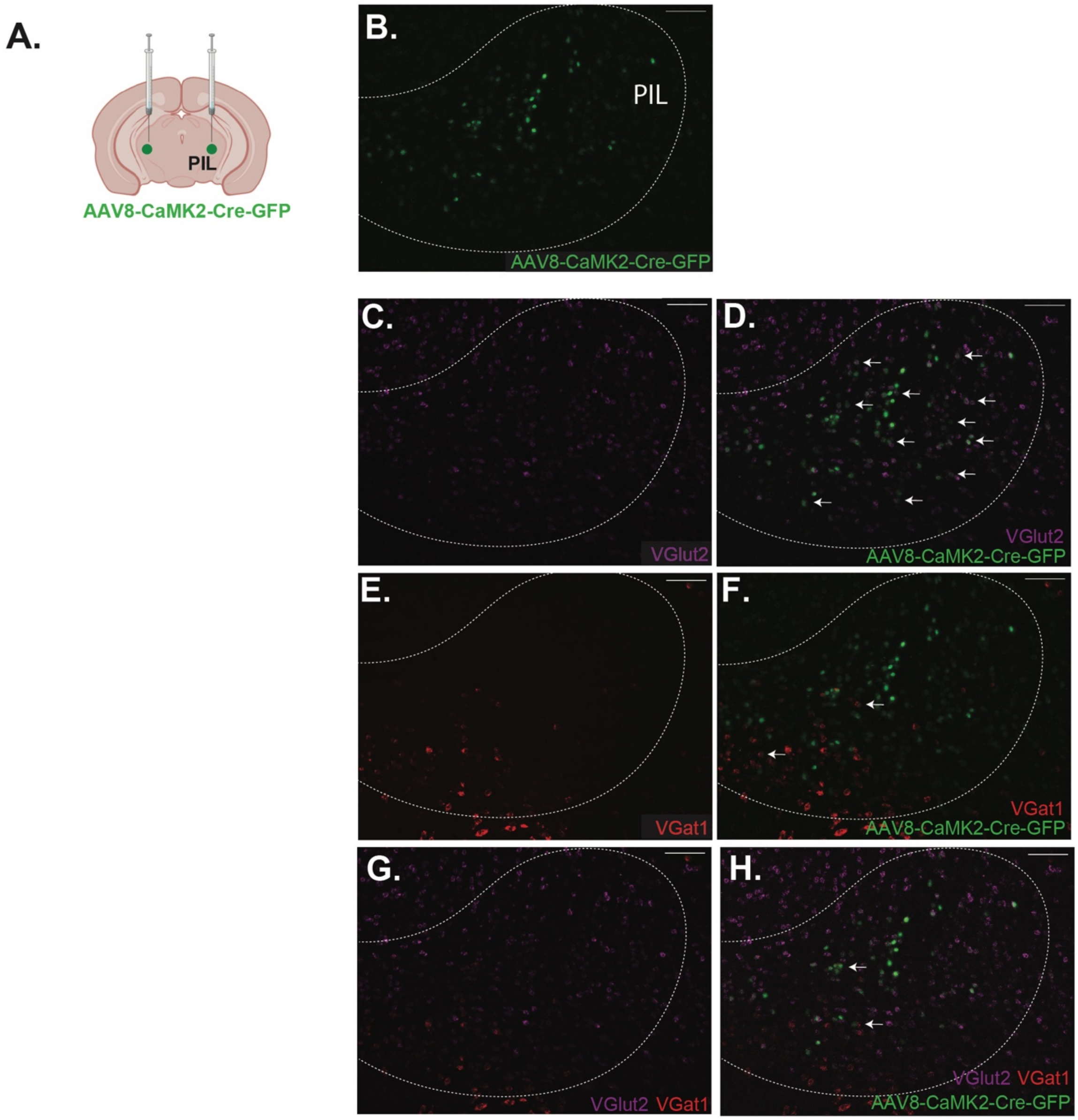

**Supplementary Figure 2. PIL neurons that express CaMKIIα are predominantly glutamatergic.** **(A)** Schematic of injection with AAV8-CaMKIIα-GFP in the PIL. **(B-H)** RNAscope performed on PIL sections containing AAV8-CaMKIIα-GFP (green; **B**) for *vglut2* (violet; **C-D**) revealed 88.82% overlap with CaMKIIα-GFP$^+$ cells. RNAscope for *vgat1* (red; **E-F**) showed 1.84% overlap with CaMKIIα-GFP$^+$ cells. 6.87% of CaMKIIα-GFP$^+$ cells co-expressed both *vglut2* and *vgat1* (**G-H**) and 2.47% expressed neither. [n = 2 males, 4 slices/subject; 20X magnification; scale bar (100 μm); arrows indicate some examples of overlap].

**Supplementary Figure 3.**

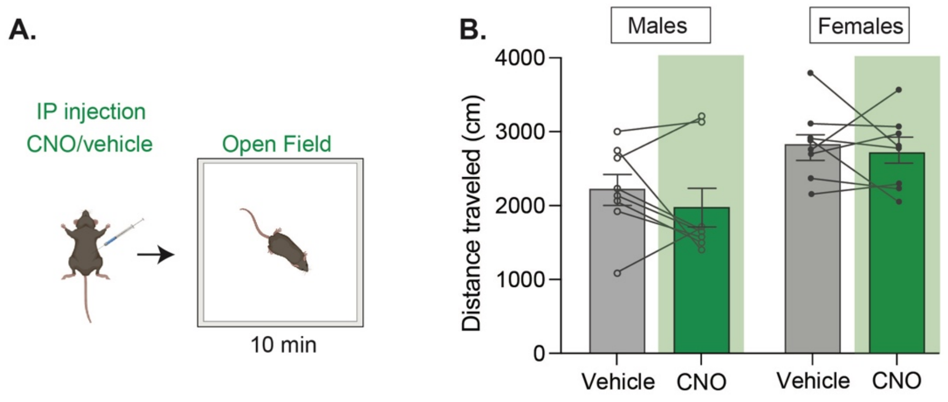


**Supplementary Figure 3. CNO administration does not impact locomotor behavior.**

**(A)** Schematic of open-field test following CNO/vehicle injection in male and female subjects without DREADD virus in the PIL. **(B)** No difference in distance traveled (centimeters) was observed following CNO or vehicle administration (Two-way mixed ANOVA; treatment $p > .05$, sex $*p < .05$, treatment x sex $p > .05$; n = 8 males and 8 females).

**Supplementary Figure 4.**

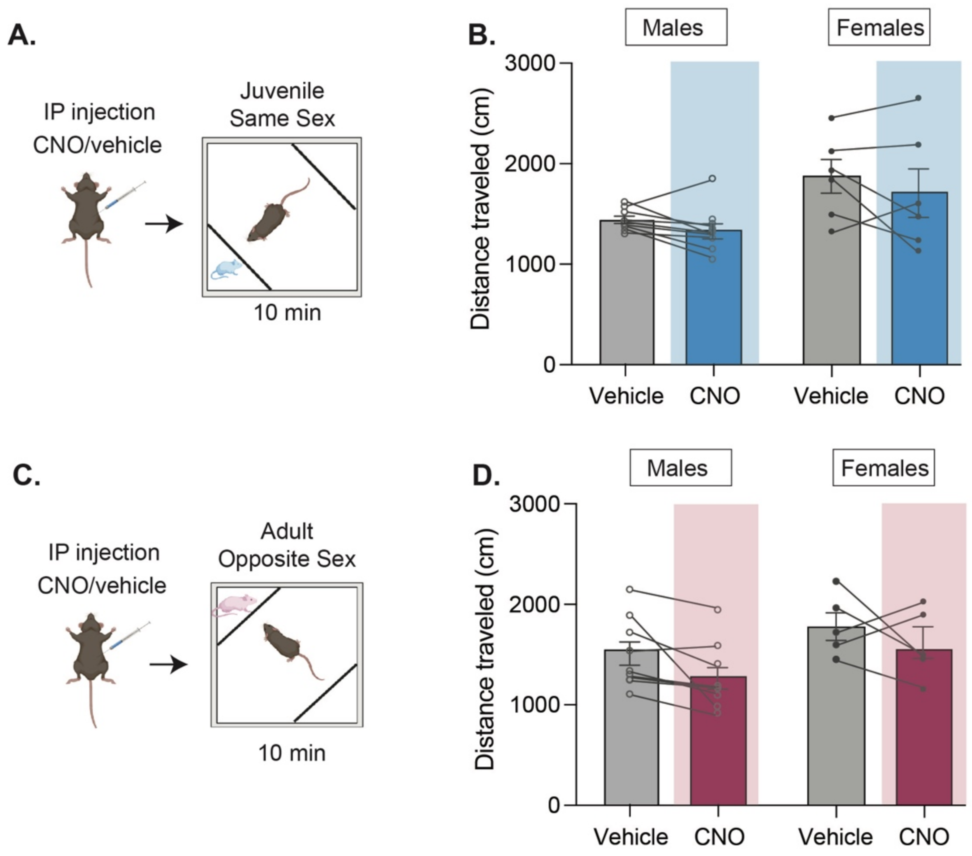


**Supplementary Figure 4. CNO administration does not impact locomotor behavior during the social preference task. (A)** Schematic of juvenile social preference task following CNO/vehicle injection in male and female subjects with CaMKIIα-GFP/hSYN-DIO-hM4D(Gi)-mCherry in the PIL. **(B)** No difference in distance traveled (cm = centimeters) was observed following CNO or vehicle administration (Two-way mixed ANOVA; treatment $p < .05$, sex $*p < .05$, treatment x sex $p > .05$; n = 9 males and 6 females). **(C)** Schematic of adult social preference task following CNO/vehicle injection. **(D)** No difference in distance traveled (cm) was observed following CNO or vehicle administration (Two-way mixed ANOVA; treatment $p > .05$, sex $p > .05$, treatment x sex $p > .05$; n = 9 males and 5 females).

**SUPPLEMENTARY REFERENCES:**

1. Netser S, Haskal S, Magalnik H, Wagner S (2017): A novel system for tracking social preference dynamics in mice reveals sex- and strain-specific characteristics. *Molecular Autism*. 8:53.

2. Cao Y, Wu R, Tai F, Zhang X, Yu P, An X, et al. (2014): Neonatal paternal deprivation impairs social recognition and alters levels of oxytocin and estrogen receptor α mRNA expression in the MeA and NAcc, and serum oxytocin in mandarin voles. *Hormones and Behavior*. 65:57-65.

3. Ferguson JN, Young LJ, Hearn EF, Matzuk MM, Insel TR, Winslow JT (2000): Social amnesia in mice lacking the oxytocin gene. *Nature Genetics*. 25:284-288.

4. Jacobs SA, Huang F, Tsien JZ, Wei W (2016): Social recognition memory test in rodents. *Bio-protocol*. 6:e1804-e1804.

5. Spiteri T, Ågmo A (2009): Ovarian hormones modulate social recognition in female rats. *Physiology & Behavior*. 98:247-250.

6. Morel C, Montgomery SE, Li L, Durand-de Cuttoli R, Teichman EM, Juarez B, et al. (2022): Midbrain projection to the basolateral amygdala encodes anxiety-like but not depression-like behaviors. *Nature Communications*. 13:1-13.

7. Martianova E, Aronson S, Proulx CD (2019): Multi-fiber photometry to record neural activity in freely-moving animals. *JoVE (Journal of Visualized Experiments)*.e60278.
